# Parasitic and Commensal interactions among Mimiviruses, Sputnik-like virophages, and Transpovirons: A theoretical and dynamical systems approach

**DOI:** 10.1101/2024.09.13.610890

**Authors:** Emmanuel Ortega-Atehortúa, Nicole Rivera Parra, Boris A. Rodríguez, Gloria Machado-Rodríguez, Juan Camilo Arboleda Rivera

## Abstract

Giant viruses have been in the scope of virologists since 2003 when they were isolated from *Acanthamoeba* spp. Giant viruses, in turn, get infected by another virus named virophage and a third biological entity that corresponds to a transpoviron which can be found in the capsids of giant and virophage viruses. So far, transpovirons seem to behave as commensal entities while some virophages exhibit commensal behavior under laboratory conditions. To study the system’s behavior, we used a theoretical approximation and developed an ordinary differential equation model. The dynamical analysis showed that the system exhibits an oscillatory robust behavior leading to a hyperparasitic Lotka-Volterra dynamic. But the biological mechanism that underlines the transpoviron persistence over time remains unclear and its status as a commensal entity needs further assessment. Also, the ecological interaction that leads to the overall coexistence of the three viral entities needs to be further studied.

## 1 Introduction

*Acanthamoeba* spp. are protists present in almost any environment. They have been studied as eukaryotic cellular motility models [1] and as pathogens causing some health issues like sinus infection [2], keratitis, and encephalitis [3]. Furthermore, they participate in soil ecology processes like nutrient cycling and microbial community structure through bacteria consumption [4]. Besides, *Acanthamoeba* sp. can act as microbial reservoirs and in some cases as trojan horses for human pathogens, such as *Helicobacter pylori, Legionella pneumophila, Listeria monocytogenes, Staphylococcus aureus, Toxoplasma gondii, Cryptosporidium* sp., *Streptomyces californicus*, and *Exophiala dermatitidis. Acanthamoeba* sp. is also subject to Acanthamoeba polyphaga mimivirus (APMV) infection, which is the largest virus known to date and belongs to the Mimiviridae family [5].

The Mimiviridae family is composed of group I viruses (double-stranded DNA) that are found within a monophyletic group called nucleocytoplasmic large DNA viruses (NCLDV) [6]. Acanthamoeba polyphaga mimivirus (APMV) is the target of another recently discovered viral entity called virophage. The first described virophage was Sputnik, which cannot replicate within Acanthamoeba polyphaga cells alone but needs the presence of APMV (Mamavirus Strain) [7]. This was further elucidated by transmission electron microscopy and polymerase chain reaction in *Acanthamoeba castellanii* coinfected with Mimivirus and Sputnik, which showed negative effects for the giant virus in the presence or the virophage instead of the target cell. These effects ranged from defective viral particles, uncontrolled membrane accumulation around capsids, DNA empty capsids, and capsids with virophage virions inside [8].

While giant viruses like mimiviruses are rather recently studied viral entities with some interesting and new biological functions, virophages might represent further complexity in terms of ecological interactions. Virophages can also enter the cellular host via clathrin- mediated endocytosis without the need for giant virus entry, and can also integrate their genomes into the cellular host [9]. This genome integration leads to a mutualistic interaction (Virophage and Acanthamoeba) and an induced altruism for the amoeba [10]. While this type of dynamics can occur for Mavirus like virophages, Sputnik like virophages seem to only integrate their genome into the Giant virus [11]. This type of virophages depend on the formation of a composite with their associated giant virus to enter the cell. This is called the paired entry mode of coinfection [12].

Transpovirons represent a new type of transposable element exclusive to giant viruses which was first detected as a high copy extrachromosomal DNA molecule and was named as a transpoviron because it is a transposable element inside a virus [9]. These transposable elements show a commensal interaction with giant viruses and virophages. Also, there is no evidence of loss in fitness related to transpoviron infection in these two viruses. This ecological interaction can also be seen in vitro conditions where the virophage Zamilon vitiis does not impair the replication cycle of Mimiviruses from the clades B and C [13].

Previous work has been done to elucidate the two types of coinfection mechanisms that virophages and giant viruses can perform, independent cell entry or simultaneous cell entry. The authors used Ordinary Differential Equations (ODE) with four state variables, growth dynamics of the cellular host, giant virus, virophage, and the virus-virophage composite [12]. However, this work does not show any of the previously mentioned ecological interactions and its possible implications for the system’s dynamics. As a result, the addition of the third viral component (Transpoviron) and the recently discovered interactions that lead to new features in this ecological network implies an opportunity to elucidate the dynamics of these interactions. The aim of this works lies in a theoretical approach to the dynamics and ecological relations amongst these three viral entities using the theory of dynamical systems.

## 2 Methods

### 2.1 Dynamic System

The mathematical model (Eq. 1 to 19) aims to represent the commensal and parasitic ecological interactions for the Acanthamoeba-Giant Virus-Virophage-Transpoviron system. The mode of infection follows the Zamilon-Sputnik-like system of coinfection [12]. ODE’s assumptions are as follows:

1. The system is well-mixed.
2. Rates of contact follow mass action kinetics.
3. All possible composite formations are independent state variables.
4. The Sputnik virophage genome does not integrate into the cell genome.
5. Whenever possible the parameter values are derived from literature.

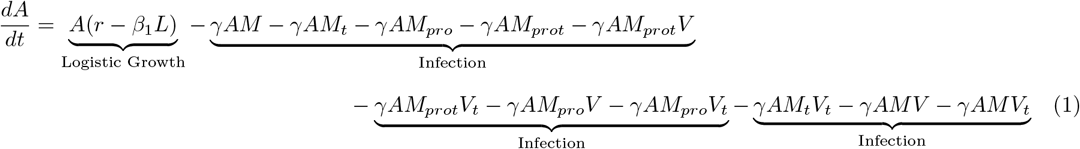

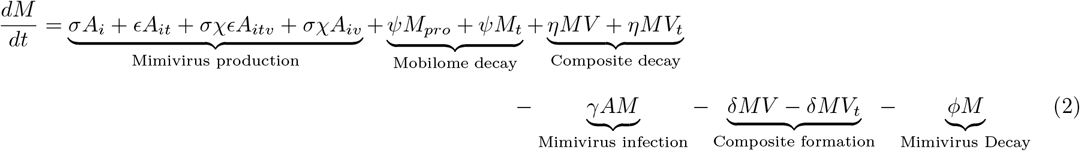

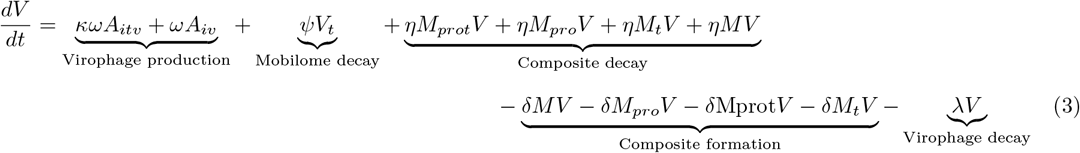

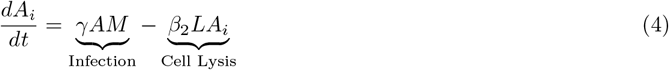

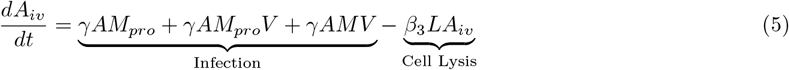

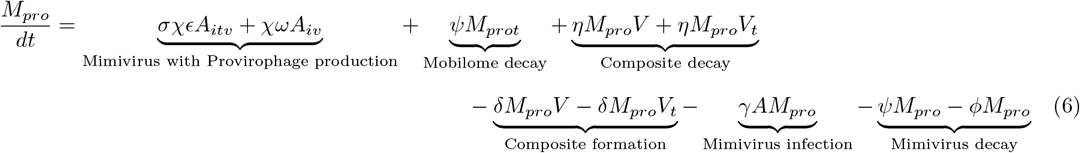

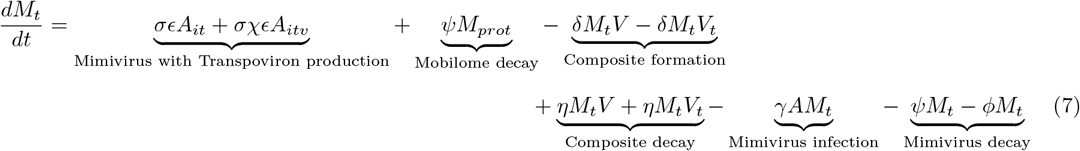

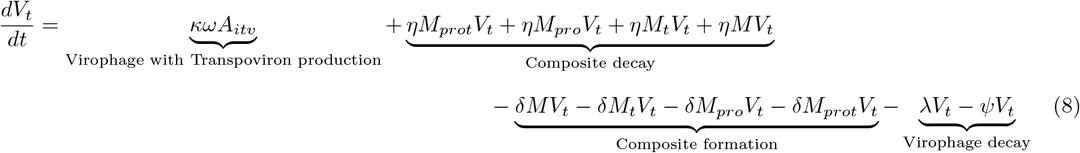

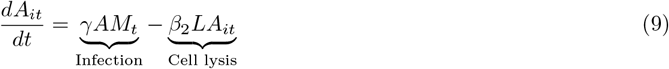

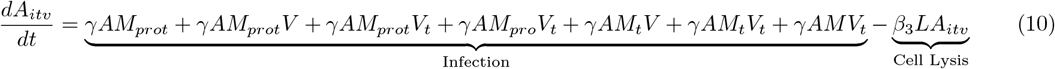

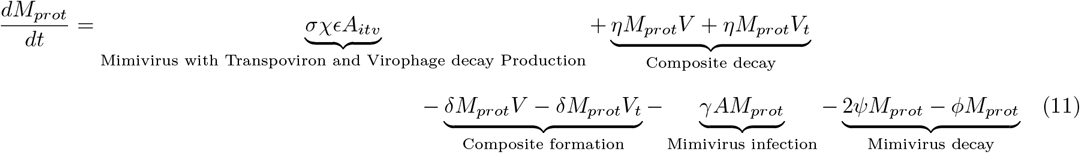

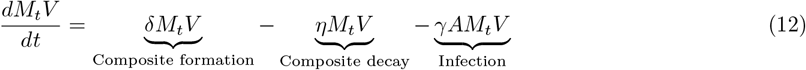

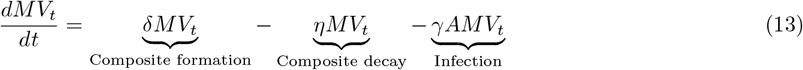

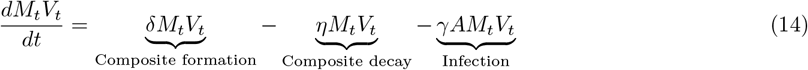

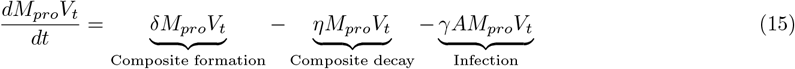

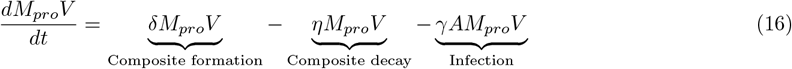

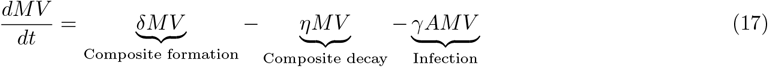

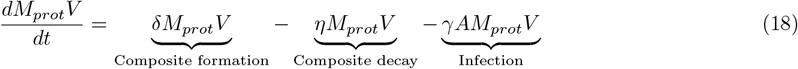

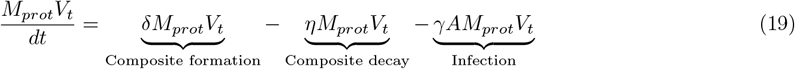

Parameters are described in Table 2.

**Table 1:** State variables and meanings. These state variables represent all the possible viral and cellular entities for the system. First the main components and second all the composite formations.

| State Variable | Meaning |
| --- | --- |
| $A$ | *Acanthamoeba* |
| $M$ | Mimivirus |
| $V$ | Virophage |
| $M_{pro}$ | Mimivirus with Provirophage |
| $A_i$ | *Acanthamoeba* infected with Mimivirus |
| $A_{iv}$ | *Acanthamoeba* infected with Mimivirus and Virophage |
| $M_t$ | Mimivirus with Transpoviron |
| $V_t$ | Virophage with Transpoviron |
| $M_{prot}$ | Mimivirus with Provirophage and Transpoviron |
| $A_{it}$ | *Acanthamoeba* infected with Mimivirus and Transpoviron |
| $A_{itv}$ | *Acanthamoeba* infected with Mimivirus, Virophage and Transpoviron |
| Composites | Viral Entities |
| $MV$ | $M$ and $V$ |
| $MV_t$ | $M$ and $V_t$ |
| $M_tV$ | $M_t$ and $V$ |
| $M_tV_t$ | $M_t$ and $V_t$ |
| $M_{pro}V$ | $M_{pro}$ and $V$ |
| $M_{pro}V_t$ | $M_{pro}$ and $V_t$ |
| $M_{prot}V$ | $M_{prot}$ and $V$ |
| $M_{prot}V_t$ | $M_{prot}$ and $V_t$ |

**Table 2:** Parameters used in the model. These parameters were all obtained from experimental data reported in literature except for Mobilome decay rate (*ψ*). The table displays the symbol, meaning, units and reference for each parameter.

| Symbol | Meaning | Value | Units | Reference |
| --- | --- | --- | --- | --- |
| $k$ | Carrying capacity | $4.0 \times 10^6$ | host/mL | [14] |
| $\alpha$ | Growth rate of non infected population | 1.4 | day<sup>-1</sup> | [14] |
| $\beta$ | Cell death rate | 0.7 | day<sup>-1</sup> | [12] |
| $\beta_2$ | Death rate of Mimivirus infected population | 0.9 | day<sup>-1</sup> | [15] |
| $\beta_3$ | Death rate of Mimivirus and Virophage infected population | 0.8 | day<sup>-1</sup> | [15] |
| $\gamma$ | Infection rate | $4.3 \times 10^{-8}$ | mL × viral particles<sup>-1</sup> × day<sup>-1</sup> | [12] |
| $\epsilon$ | Composite formation rate | $2.2 \times 10^{-8}$ | mL × viral particles<sup>-1</sup> × day<sup>-1</sup> | [12] |
| $\phi$ | Mimivirus death rate | $3.2 \times 10^{-2}$ | day<sup>-1</sup> | [16] |
| $\lambda$ | Virophage death rate | $3.2 \times 10^{-1}$ | day<sup>-1</sup> | [12] |
| $\psi$ | Mobilome decay rate | 0.001 | day<sup>-1</sup> | Assumed |
| $\chi$ | Control Parameter (Virophage to Mimivirus) | 0.3 | dimensionless | [15] |
| $\epsilon$ | Control Parameter (Transpoviron to Mimivirus) | 1 | dimensionless | [13] |
| $\kappa$ | Control Parameter (Transpoviron to Virophage) | 1 | dimensionless | [13] |
| $\eta$ | Composite decay rate | $(\lambda + \phi)$ | day<sup>-1</sup> | [12] |
| $\zeta$ | Composite outbreak | $30 * \beta_3$ | viral particles/host | [12] |
| $\sigma$ | Mimivirus outbreak | $300 * \beta_2$ | viral particles/host | [17] |
| $\omega$ | Virophage outbreak | $1500 * \beta_3$ | viral particles/host | [12] |

For the initial parameter values the Transpoviron was assumed to behave as a commensal entity to the Mimivirus and the Virophage [13], while the virophage started as a parasitic entity to the Mimivirus. The three control parameters and its base values are *χ* = 0.3 (Virophage inhibition to Mimivirus), *ϵ* = 1 (Transpoviron inhibition to Mimivirus), *κ* = 1 (Transpoviron inhibition to Virophage). Where 0 means total inhibition (Parasitism) and 1 means no inhibition (Commensalism).

### 2.2 Computational Methods

Simulations were performed in Wolfram Mathematica 13 [18] and Python 3 [19, 20, 21]. The ODE system was computed using the numerical solver `NDSolve` with the `StiffnessSwitching` Method. The Steady state of the system and the linear stability analysis of the eigenvalues were also calculated in Mathematica.

### 2.3 Linear Stability Analysis

First the fixed points of the ODE were obtained, and the Jacobian matrix was constructed. The linear stability analysis was performed by using the function `EigenValues` in Mathematica to determine the behavior around the fixed points for the base values and standard conditions of the system [22].

### 2.4 Global Sensitivity and Bifurcation Analysis

To perform the bifurcation analysis, the fixed points of the system were calculated 1331 times. Each calculation had different combinations of values for the three control parameters (*χ, ϵ, κ*) that ranged from 0 to 1 with a granularity of 0.1. Finally, the bifurcation diagrams were plotted and where changes of the fixed-point values were detected, the derivative was calculated to determine the global sensitivity [23].

### 2.5 Local Sensitivity Analysis

Sensitivity analysis was performed for the same parameters as in the bifurcation analysis. The changes in the parameter values were 1%, 3% and 5% deviations from the base values to verify the accuracy of the finite difference approach [24]. Then the fixed points were calculated for each percentual change in value for each parameter. For each parameter the Euclidean distance was calculated to make sure the compared fixed points between each set of parameter changes were the same. Finally, the Absolute Sensitivity Coefficient was calculated (Eqn 20).

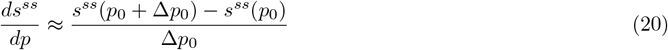

### 2.6 Data availability

The code is available at https://github.com/Mimivirus16/Transpoviron-ODE.

### 3 Results

### 3.1 Transpoviron Coexistence

Stable coexistence for the four biological entities does not show for base parameter values and literature initial conditions as shown in Figure 1A. This suggests that the transpoviron would become extinct over time in these conditions. However, this type of dynamic does not seem possible due to the co-existing dynamics that the four biological entities have shown in some experiments. As previously stated by Jeudy et al. [13], the transpoviron is a commensal viral entity because it does not affect the fitness of the Mimivirus and Virophage. Even so, this ecological interaction for this system can be further assessed by theoretical and mathematical approaches.

**Figure 1:**
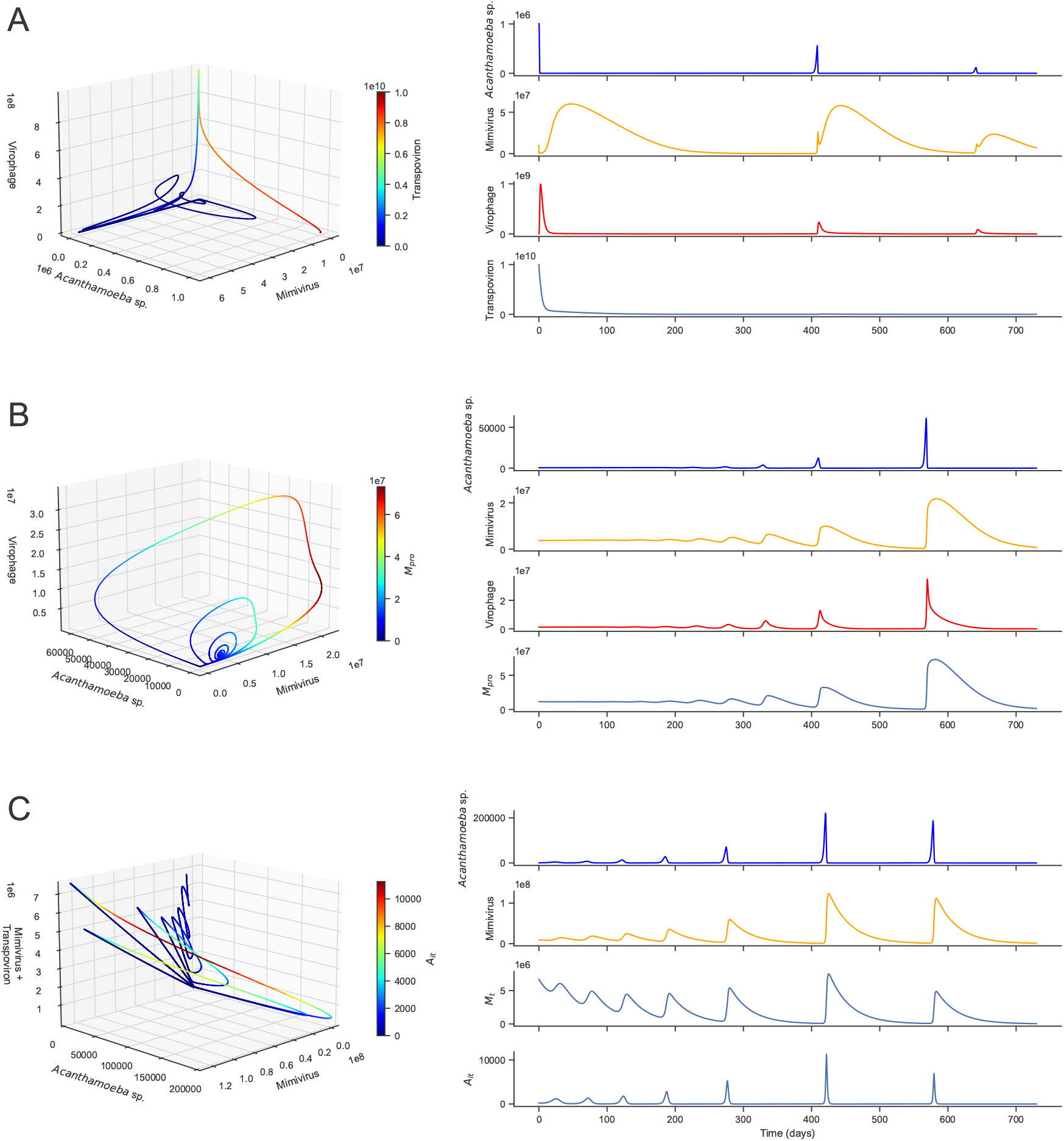
Phase Diagrams. *In silico* simulations for the next conditions: A) Real parameter values with experimental initial conditions: *A* = 1 *×* 10^6^, *M* = 10 *×* 10^6^, *V* = 0, *A*_*i*_ = 0, *A*_*iv*_ = 0, *M*_*pro*_ = 0, *M*_*t*_ = 0, *V*_*t*_ = 10 *×* 10^9^, *A*_*it*_ = 0, *A*_*itv*_ = 0, *M*_*prot*_ = 0, *M*_*t*_*V* = 0, *MV*_*t*_ = 0, *M*_*t*_*V*_*t*_ = 0, *M*_*pro*_*V*_*t*_ = 0, *M*_*pro*_*V* = 0, *MV* = 0, *M*_*prot*_*V* = 0, *M*_*prot*_*V*_*t*_ = 0 [13]; the simulation time was 730 days. B) In this simulation that represents the commensalism scenario, the parameter values were equal to A, except *χ* = 1; initial conditions were: *A* = 465.478, *M* = 3.71043 *×* 10^6^, *V* = 1.1811 *×* 10^6^, *A*_*i*_ = 82.6775, *A*_*iv*_ = 314.98, *M*_*pro*_ = 1.14463 *×* 10^7^, *Mt* = 0, *V*_*t*_ = 0, *A*_*it*_ = 0, *A*_*itv*_ = 0, *M*_*prot*_ = 0, *M*_*t*_*V* = 0, *MV*_*t*_ = 0, *M*_*t*_*V*_*t*_ = 0, *M*_*pro*_*V*_*t*_ = 0, *M*_*pro*_*V* = 844908, *MV* = 273885, *M*_*prot*_*V* = 0, *M*_*prot*_*V*_*t*_ = 0; This was obtained from Bifurcation Fixed point Number 5841 (Table S1). C) In this simulation that exhibits the stable equilibrium for the Transpoviron, the parameter values were equal to A except *κ* = 0.99. Initial conditions were obtained from the 1% Sensitivy Analysis: *A* = 640.251, *M* = 9.58125 *×* 10^6^, *V* = 0, *A*_*i*_ = 293.005, *A*_*iv*_ = 0, *M*_*pro*_ = 0, *M*_*t*_ = 6.69319 *×* 10^6^, *V*_*t*_ = 0, *A*_*it*_ = 204.685, *A*_*itv*_ = 0, *M*_*prot*_ = 0, *M*_*t*_*V* = 0, *MV*_*t*_ = 0, *M*_*t*_*V*_*t*_ = 0, *M*_*pro*_*V*_*t*_ = 0, *M*_*pro*_*V* = 0, *MV* = 0, *M*_*prot*_*V* = 0, *M*_*prot*_*V*_*t*_ = 0.

### 3.2 Bifurcation analysis reveals system coherence and robust qualitative Dynamics

The overall possible dynamics for the three-parameter space are as shown in Figure 2A. With a granularity of 0.1 the transpoviron did not appear in the continuum of the bifurcation analysis. Furthermore, the stability of all the fixed points did not change over time, showing the robust and stable behavior of the system. Also, all the stable equilibria for which the Virophage appeared were always accompanied by the Provirophage state in the Mimivirus genome suggesting that the Virophage would only appear together with the Mimivirus if the genome integration occurs. The dynamics for this type of fixed-points are shown in Figure 1B. Moreover, the bifurcation diagrams in Figure 2B shows a bifurcation in *χ* = 0.2 for state variables *A, M* and *V*. Fixed Points below this value do not show a stable equilibrium where the virophage coexists, suggesting that a parasitoid behavior for this entity makes it a non-stable population. The derivatives associated with bifurcation point AMPV in Table 3 showed that The *χ* derivative associated with the Virophage is greater than that of the Mimivirus only when *χ >* 0.46.

**Table 3:** Derivatives associated to the AMPV fixed point for state variables *A, M* and *V. s*^*ss*^ refers to the value of a given state variable in the steady state.

| State variable | $\frac{ds^{ss}}{d\chi}$ |
| --- | --- |
| $A$ | $\frac{-448.481}{\chi^{1.7}}$ |
| $M$ | $\frac{-286.816}{\chi^{2.8}}$ |
| $V$ | $\frac{-1.51749 \times 10^6}{\chi}$ |

**Figure 2:**
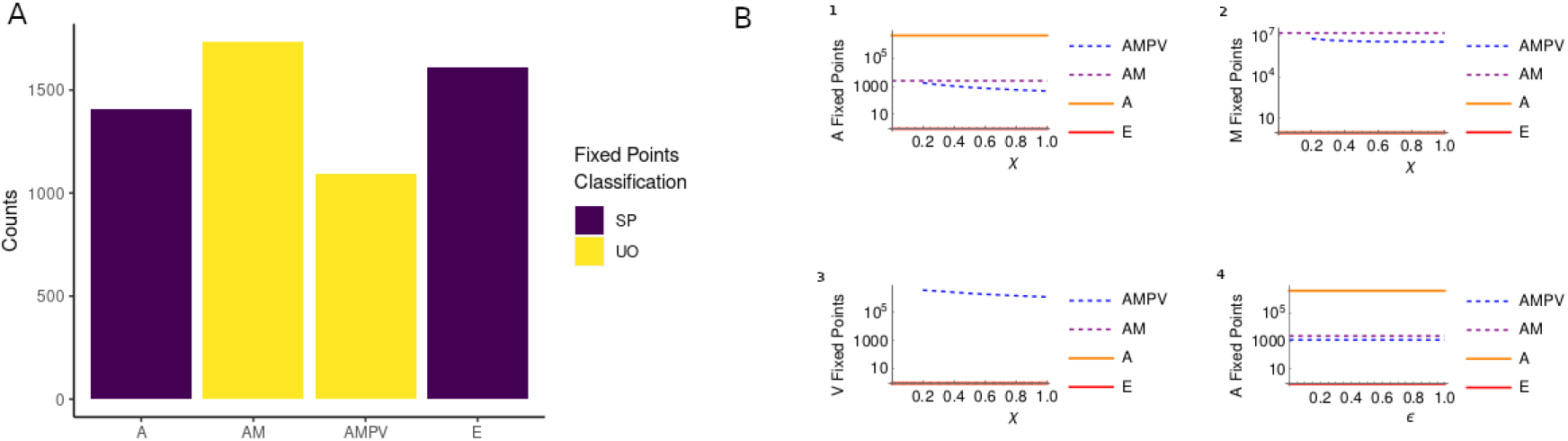
Bifurcation Analysis. This figure shows the results from the bifurcation analysis for the 1331 combinations for parameters *χ, ϵ* and, *κ*. A=Bifurcation Analysis Histogram, the Y axis shows the number of times the fixed point was present. The X axis shows the coexisting populations in a fixed point where A: *Acanthamoeba* sp. alone, AM: *Acanthamoeba* sp. and Mimivirus, AMPV: *Acanthamoeba* sp., Mimivirus (Provirophage) and Virophage, E: Extinction of all populations. The fixed-point classifications are shown in the right side of the graphic where SP=Saddle Point and UO=Unstable Oscillator. B=Bifurcation diagrams, the Y axis for each diagram shows the value of the fixed point for each value of *χ* (1,2,3) and *ϵ* for bifurcation diagram 4.

### 3.3 The Finite Approximation is not a Robust Analysis for this kind of Systems

The sensitivity analysis was performed as proposed by Ingalls [24]. However, the finite approximation approach showed inconsistencies for the fixed-point value for each state variable between all the deviations of the parameters (1%, 3% and 5%). This suggests that this approach does not work as a measure of the sensitivity of a high dimensional system. Even so, some interesting results appeared for such small changes of the parameter values. All infinitesimal changes in the parameter values showed bifurcations, suggesting a high sensitivity for this system to the change in the three control parameters (*χ, ϵ, κ*).

### 3.4 Transpoviron is not entirely commensal to the Virophage

The changes in the parameters previously calculated showed that for every infinitesimal change in *κ* a new fixed point aroused showing the coexistence of the Transpoviron with the Mimivirus. This coexistence is shown in Figure 1C. This coexistence shows that a small negative effect in the Virophage by the Transpoviron leads to a stable equilibrium and non-extinction of the Transpoviron.

## 4 Discussion

The Mimivirus-Virophage-Transpoviron hyperparasitic system emerges as a complex ecological interaction at the molecular scale that is poorly understood. However, the system presented here aroused results that indicate deeply that the transpoviron and its biological properties are poorly studied or that some of its intrinsic characteristics have been overlooked. Experimentally the transpoviron persists as part of the mobilome of Mimiviruses and Sputnik Virophages [9]. Nevertheless, the transpoviron here modeled as a transposon that inserts itself into the Mimivirus and Virophage genome could not persist over time when simulating the literature based initial conditions (Figure 1A). On top of that, when exploring the bifurcation parameter sampling, there is no fixed point where the transpoviron exists as a stable population over time (Fig 2A). The granularity used for the bifurcation analysis and the base values for the control parameters showed that for those specific values the transpoviron is not a stable population.

Furthermore, the literature focuses on the transpoviron as a commensal entity based on *in vitro* conditions to the Mimivirus but overlooks the ecological interaction that the transpoviron could show to the virophage [13]. This overlooking was addressed in the construction of the model as the control parameter that stablishes the transpoviron to virophage interaction was included (*κ*). The local sensitivity analysis showed that for small changes in *κ* (1%, 3%, 5%) fixed points where the transpoviron behaves as a stable population overtime where obtained (Fig 1C). This means that the transpoviron could persist as a stable population if it has a small cost to the virophage fitness. However, the transpoviron persistence shown in those fixed points excluded the virophage totally (AMT=Acanthamoeba, Mimivirus and transpoviron). Also, these types of interactions exist in other virophage systems like the Mavirus virophages integrated into the genome of the flagellate, *Cafeteria burkhardae*. These experiments showed that during Giant virus infection, some retrotransposons were excised from EMALES (Endogenous Mavirus Like Elements) genome. It was also demonstrated in these experiments that some non-LTR retrotransposons of the Ngaro superfamily inside the EMALES sequences were able to interrupt the provirophage expression. This shows that provirophages and retrotransposon elements can have more than just commensalism interactions leading to parasitic interactions like inactivation or excision [25].

The bifurcation analysis showed that the qualitative dynamics of the system are robust. The fixed points A, E and AM appeared over all the bifurcation continuum and their stability did not change for all 1331 combinations of control parameter values (*χ,ϵ, κ*). The only parameter that exhibited a bifurcation was the parameter that controls the virophage to mimivirus interaction *χ* (Figure 2B). This revealed that for the virophage to persist over time a parasitoid behavior is not viable *χ <* 0.2. The new fixed point for *χ >* 0.2 was AMPV, this type of dynamic was the only present as no AMV (Acanthamoeba, Mimivirus and Virophage) dynamics were found. This demonstrates that for the Virophage to persist as a stable population not only cannot behave as a parasitoid entity but also its provirophage state would persist. This is consistent with the findings of another theoretical model which also shows that the virophage will persist in the mimivirus genome as long as it does not completely affects the giant virus replication [26]. Additionally, we found that for AMPV fixed point, the virophage showed a greater sensitivity for *χ* when *χ* ≤ 0.396 compared to the mimivirus shown in Table 3.

The overall total coexistence of Acanthamoeba-Mimivirus-Virophage-Transpoviron (AMTV) in any of its possible states did not appear in any of the parameter combinations whether it was bifurcation or sensitivity analysis, thus, leaving the system with equilibrium points where A, AM, AMPV, AMT and E are possible. In the end, showing that the ecological interactions between these three viral entities depend on other not studied variables that integrate the full scope of this system. This case could be compared to the simple model of hyperparasitism for hypoviruses that infect *Cryphonectria parasitica*, the fungus responsible for chestnut blight. This system shows that high vertical transmission rates between fungi and low virulence from the hyperparasite lead to better establishment in the population [27]. However, this type of system is highly different because Mimiviruses and Virophages do not transmit the transpoviron in the same way as a fungus would transmit its viral parasite. Because the hypovirus needs the fungus to reproduce or to fuse with other fungus to spread, as most hypovirus are capsidless and don’t infect fungi as free particles [28], while Mimivirus and Virophages are free particles in the environment that together could infect the *Acanthamoeba* and the Transpoviron has two options to insert its sequence, thus leading to a more complex network and different needs to spread and replicate. Also, oscillatory and classical Lotka-Volterra dynamics were not observed for this simplified model, while the model here proposed and previous models show the classical predatory-prey oscillations [12]. Other theoretical models for the sputnik-like virophages which insert their genome into the cellular host genome also showed that this is not a stable mechanism for sputnik like viruses which is congruent to both experimental data and the dynamics proposed for this system [26].

The model here presented offers a high dimensional system that exhibits the dynamics of hypeparasistism at the cellular-molecular scale, showing the propagation of different viral entities including the transpoviron. This system questions the assumption of the transpoviron as a commensal entity and the need for further assessment of the mechanism of persistence of the transpoviron within this system as its only way of propagation is through the insertion into either the Mimivirus or Virophage genome, leading to a need of experimental verification of its overall replication mechanism.

## References

[1] T. D. Pollard, E. Shelton, R. R. Weihing, and E. D. Korn, “Ultrastructural characterization of f-actin isolated from acanthamoeba castellanii and identification of cytoplasmic filaments as f-actin by reaction with rabbit heavy meromyosin,” Journal of Molecular Biology, vol. 50, pp. 91–IN24, 5 1970.

[2] G. S. Visvesvara, H. Moura, and F. L. Schuster, “Pathogenic and opportunistic free-living amoebae: Acanthamoeba spp., balamuthia mandrillaris, naegleria fowleri, and sappinia diploidea,” FEMS Immunology Medical Microbiology, vol. 50, pp. 1–26, 6 2007.

[3] N. A. Khan, “Acanthamoeba : biology and increasing importance in human health,” FEMS Microbiology Reviews, vol. 30, pp. 564–595, 7 2006.

[4] J. L. Sinclair, J. F. McClellan, and D. C. Coleman, “Nitrogen mineralization by acanthamoeba polyphaga in grazed pseudomonas paucimobilis populations,” Applied and Environmental Microbiology, vol. 42, p. 667, 10 1981.

[5] R. Siddiqui and N. A. Khan, “Biology and pathogenesis of acanthamoeba,” Parasites vectors, vol. 5, 2012.

[6] B. L. Scola, S. Audic, C. Robert, L. Jungang, X. de Lamballerie, M. Drancourt, R. Birtles, J.-M. Claverie, and D. Raoult, “A giant virus in amoebae,” Science, vol. 299, pp. 2033–2033, 3 2003.

[7] R. Smallridge, “A virus gets a virus,” Nature Reviews Microbiology, vol. 6, p. 714, 2008.

[8] B. L. Scola, C. Desnues, I. Pagnier, C. Robert, L. Barrassi, G. Fournous, M. Merchat, M. Suzan-Monti, P. Forterre, E. Koonin, and D. Raoult, “The virophage as a unique parasite of the giant mimivirus,” Nature 2008 455:7209, vol. 455, pp. 100–104, 8 2008.

[9] C. Desnues, B. L. Scola, N. Yutin, G. Fournous, C. Robert, S. Azza, P. Jardot, S. Monteil, A. Campocasso, E. V. Koonin, and D. Raoult, “Provirophages and transpovirons as the diverse mobilome of giant viruses,” Proceedings of the National Academy of Sciences, vol. 109, pp. 18078–18083, 10 2012.

[10] E. V. Koonin and M. Krupovic, “Polintons, virophages and transpovirons: a tangled web linking viruses, transposons and immunity,” Current Opinion in Virology, vol. 25, pp. 7–15, 8 2017.

[11] C. Desnues and D. Raoult, “Inside the lifestyle of the virophage,” Intervirology, vol. 53, pp. 293–303, 6 2010.

[12] B. P. Taylor, M. H. Cortez, and J. S. Weitz, “The virus of my virus is my friend: Ecological effects of virophage with alternative modes of coinfection,” Journal of Theoretical Biology, vol. 354, pp. 124–136, 8 2014.

[13] S. Jeudy, L. Bertaux, J.-M. Alempic, A. Lartigue, M. Legendre, L. Belmudes, S. Santini, N. Philippe, L. Beucher, E. G. Biondi, S. Juul, Daniel, J. Turner, Y. Couté, J.-M. Claverie, and C. Abergel, “Exploration of the propagation of transpovirons within mimiviridae reveals a unique example of commensalism in the viral world,” The ISME Journal, vol. 14, pp. 727–739, 2020.

[14] T. J. Byers, R. A. Akins, B. J. Maynard, R. A. Lefken, and S. M. Martin, “Rapid growth of acanthamoeba in defined media; induction of encystment by glucose-acetate starvation,” The Journal of protozoology, vol. 27, pp. 216–219, 1980.

[15] B. Tokarz-Deptula, S. Chrzanowska, N. Gurgacz, M. Stosik, and W. Deptula, “Virophages—known and unknown facts,” Viruses 2023, Vol. 15, Page 1321, vol. 15, p. 1321, 6 2023.

[16] R. K. Campos, K. R. Andrade, P. C. P. Ferreira, C. A. Bonjardim, B. L. Scola, E. G. Kroon, and J. S. Abrahão, “Virucidal activity of chemical biocides against mimivirus, a putative pneumonia agent,” Journal of Clinical Virology, vol. 55, pp. 323–328, 12 2012.

[17] V. E. J., Lesser Known Large dsDNA Viruses, vol. 328 pp. 89–121. Springer Berlin Heidelberg, 2009.

[18] W. R. Inc., “Mathematica, Version 13.3.” Champaign, IL, 2023.

[19] P. Virtanen, R. Gommers, T. E. Oliphant, M. Haberland, T. Reddy, D. Cournapeau, E. Burovski, P. Peterson, W. Weckesser, J. Bright, S. J. van der Walt, M. Brett, J. Wilson, K. J. Millman, N. Mayorov, A. R. J. Nelson, E. Jones, R. Kern, E. Larson, C. J. Carey, İ. Polat, Y. Feng, E. W. Moore, J. VanderPlas, D. Laxalde, J. Perktold, R. Cimrman, I. Henriksen, E. A. Quintero, C. R. Harris, A. M. Archibald, A. H. Ribeiro, F. Pedregosa, P. van Mulbregt, and SciPy 1.0 Contributors, “SciPy 1.0: Fundamental Algorithms for Scientific Computing in Python,” Nature Methods, vol. 17, pp. 261–272, 2020.

[20] C. R. Harris, K. J. Millman, S. J. van der Walt, R. Gommers, P. Virtanen, D. Cournapeau, E. Wieser, J. Taylor, S. Berg, N. J. Smith, R. Kern, M. Picus, S. Hoyer, M. H. van Kerkwijk, M. Brett, A. Haldane, J. Fernández del Río, M. Wiebe, P. Peterson, P. Gérard-Marchant, K. Sheppard, T. Reddy, W. Weckesser, H. Abbasi, C. Gohlke, and T. E. Oliphant, “Array programming with NumPy,” Nature, vol. 585, p. 357–362, 2020.

[21] J. D. Hunter, “Matplotlib: A 2d graphics environment,” Computing in Science & Engineering, vol. 9, no. 3, pp. 90–95, 2007.

[22] S. H. Strogatz, Nonlinear Dynamics and Chaos. CRC Press, 2 ed., 5 2015.

[23] A. Saltelli, M. Ratto, T. Andres, F. Campolongo, J. Cariboni, D. Gatelli, M. Saisana, and S. Tarantola, Global Sensitivity Analysis. The Primer. Wiley, 12 2007.

[24] B. Ingalls, Mathematical Modeling in Systems Biology. 1 2013.

[25] A. Koslová, T. Hackl, F. Bade, A. Sanchez Kasikovic, K. Barenhoff, F. Schimm, U. Mersdorf, and M. G. Fischer, “Endogenous virophages are active and mitigate giant virus infection in the marine protist Cafeteria burkhardae,” vol. 121, no. 11, p. e2314606121.

[26] J. G. Nino Barreat and A. Katzourakis, “Ecological and evolutionary dynamics of cell-virus-virophage systems,” PLOS Computational Biology, vol. 20, pp. 1–23, 02 2024.

[27] A. Y. Morozov, C. Robin, and A. Franc, “A simple model for the dynamics of a host–parasite–hyperparasite interaction,” Journal of Theoretical Biology, vol. 249, pp. 246–253, 11 2007.

[28] M. N. Pearson, R. E. Beever, B. Boine, and K. Arthur, “Mycoviruses of filamentous fungi and their relevance to plant pathology,” Molecular Plant Pathology, vol. 10, p. 115, 1 2009.

